# scTransMIL Bridges Patient-Level Phenotypes and Single-Cell Transcriptomics for Cancer Screening and Heterogeneity Inference

**DOI:** 10.1101/2025.04.22.649948

**Authors:** Zhenchao Tang, Fang Wang, Fan Yang, Jiangning Song, Calvin Yu-Chian Chen, Jianhua Yao

**Affiliations:** School of AI for Science, Peking University, Beijing, China; AI for Life Science Lab, Tencent, Shenzhen, China; School of Intelligent Systems Engineering, Shenzhen Campus of Sun Yat-sen University, Shenzhen, China; Biomedicine Discovery Institute and Department of Biochemistry and Molecular Biology, Monash University, Melbourne, Australia; State Key Laboratory of Chemical Oncogenomics, Key Laboratory of Chemical Genomics, School of Chemical Biology and Biotechnology, Peking University Shenzhen Graduate School, Shenzhen, China; Department of Medical Research, China Medical University Hospital, Taichung, Taiwan

**Author notes:** Correspondence to: Fang Wang, Calvin Yu-Chian Chen, Jianhua Yao. Equal contribution (Zhenchao Tang, Fang Wang).

## Abstract

Single-cell sequencing technology has revolutionized cancer research by revealing unprecedented insights into tumor heterogeneity. However, reliably connecting patient-level cancer phenotypes with single-cell transcriptomic profiles remains challenging, due to technical constraints and labeling ambiguity. This further hinders the precise cancer screening and intensive study of tumor mechanism based on single-cell sequencing. To bridge this gap, we introduce scTransMIL, a **sc**RNA-seq **Trans**former-based **M**ulti-**I**nstance **L**earning framework that learns whole-genome context to deliver comprehensive cancer insights at the sample, cell, and gene levels across three biological scales: (1) accurate patient-level cancer phenotype prediction, (2) precise single-cell disease scoring (validated on 4 million single cells), and (3) genome-wide biomarker discovery. Benchmark experiments demonstrate scTransMIL’s exceptional performance, including robust out-of-distribution generalization and clinically relevant prediction of metastatic tissue-of-origin, a crucial capability for identifying cancers of unknow primary. At single-cell resolution, scTransMIL identified both known and novel biomarkers for tumor B cells that conventional differential expression analysis failed to detect while maintaining consistent concordance with malignant cell annotations. scTransMIL’s adaptability is exemplified in acute myelocytic leukemia, where minimal fine-tuning with only a few patient sample labels enabled: (i) discovery of novel disease subtypes with distinct clinical outcomes, (ii) reconstruction of differentiation trajectories at the single-cell resolution, and (iii) identification of subtype-specific gene signatures. In summary, by systematically linking cellular and molecular profiles with clinical disease phenotypes, scTransMIL emerges as a transformative tool poised to advance both basic cancer research and precision oncology applications.

## Introduction

Single-cell sequencing technology is revolutionizing cancer research and clinical diagnosis by providing unprecedented resolution into tumor heterogeneity and disease progression^1^. This high-resolution approach enables early cancer detection and improved patient outcomes^2^, while also revealing the complex genomic diversity that underlies variations in cancer subtypes and treatment responses. The ability to deconstruct patient-specific tumor cell populations and their molecular profiles is critical for developing personalized and targeted treatment strategies^3^. As single-cell RNA sequencing (scRNA-seq) becomes increasingly widespread and datasets grow exponentially, artificial intelligence (AI) has been widely adopted for fundamental scRNA-seq analysis tasks^4–6^ including imputation^7–12^, integration^13–19^, and cell annotation^20–26^. However, AI applications that directly connect scRNA-seq data to disease phenotypes remains underdeveloped^27^, hampered by significant technical challenges. Current approaches for labeling disease-associated single cells predominantly rely on copy-number variant (CNV) inference, which is limited to genetically unstable cells^28^ and often confounded by the presence of normal cells in tumor samples that obscure CNV signals, especially when analyzing whole-genome data^29,30^.

Publicly available cancer-related scRNA-seq datasets typically contain valuable patient-level phenotype annotations, but the lack of single-cell resolution labels restricts their utility for understanding cancer biology at the cellular level^31–33^. To address this limitation, computational methods that are capable of bridging patient-level phenotypes with single-cell resolution insights are needed. Against the backdrop of rapid advancements in AI, we aim to develop a data-driven deep learning model that achieves the three key objectives: accurately predicting cancer phenotypes of a given patient sample for early diagnosis, inferring cancer-relatedness at single-cell resolution to uncover underlying cellular mechanisms driving the cancer phenotype, and identifying tumor-specific genes across the whole genome to facilitate biomarker discovery and precision oncology.

Current approaches for linking patient-level cancer phenotypes to single-cell data can be categorized into three main types, each with notable limitations: i) The first, supervised learning on pseudo-bulk data (Methods like KIDA), aggregates single-cell data into pseudo-samples for sample-level classification but sacrifices single-cell resolution, making it difficult to capture cellular heterogeneity^34^; ii) The second, reference mapping via bulk and single-cell data integration^35^ (e.g., DEGAS), aligns bulk and single-cell data in a shared latent space, yet excessive focus on data alignment often diminishes the heterogeneity of the latent representation, weakening the methods’ phenotype prediction accuracy^36^; iii) The third, weakly-supervised learning (e.g., scIDST), leverages patient-level disease phenotype labels to estimate disease-relatedness at the single-cell resolution^37,38^ but tends to prioritize easily detectable positive cells, overlooking subtler yet biologically relevant cell populations. Additionally, in terms of interpretability, both the second and third types of methods take the dimension-reduced cell representations as inputs, which prevents the model from directly mapping predictions back to gene-level insights^39^.

To address these limitations, we introduce scTransMIL, a **sc**RNA-seq **Trans**former-based **M**ulti-**I**nstance **L**earning (MIL) framework designed to bridge patient-level cancer phenotypes and single-cell data. MIL, a weakly supervised learning approach widely used in whole-slide image analysis, has demonstrated its effectiveness in handling weakly labeled biomedical data^40–45^. It treats patient samples as “bags”, and individual cells within samples as “instances”. However, conventional MIL approaches suffer from critical shortcomings: Instance-based MIL inaccurately assigns the bag’s label to all cells, introducing noise, while bag-based MIL optimizes only at the sample level, often over-fitting to easily identifiable positive cells while neglecting harder-to-classify yet disease-relevant instances^46^. scTransMIL addresses these issues through three key innovations. First, it integrates contrastive learning to enhance the discrimination between positive and negative cells, improving the identification of challenging yet biologically meaningful cell populations. Second, its transformer-based architecture learns genome-wide contextualized cell representations. Third, self-attention mechanisms enabling direct gene-level cancer phenotype-specific interpretability.

Trained on large-scale pan-cancer datasets, scTransMIL demonstrates superior performance in predicting cancer phenotypes for unseen patient samples and inferring cancer-relatedness at single-cell resolution compared to existing methods. Moreover, scTransMIL provides gene-level insights into tumor biology and exhibits robust generalization across independent cancer datasets. These capabilities position scTransMIL as a powerful tool for advancing cancer diagnosis, dissecting tumor heterogeneity, and accelerating biomarker discovery.

## Results

### Overall model architecture of scTransMIL

In the scTransMIL framework, MIL treats each patient sample as a “bag”, and individual cells within the sample as “instances”. While cancer phenotypes (bag labels) are available at the patient level, the phenotype of each cell (instance label) remains unknown. According to the theoretical assumption of MIL, a bag is labeled as normal only if all its instances are normal cells; conversely, any bag containing at least one tumor-related cell is labeled as a tumor bag (**Figure 1a**). Our MIL framework relies solely on bag labels for supervised learning. When processing a new bag, the model predicts both the cancer phenotype of the bag, and at the same time, quantifies the cancer-relatedness of each cell within it. As shown in **Figure 1b**, the model not only predicts whether the query bag is a tumor bag, but also infers the disease-related score of all instances within this bag.

**Figure 1.**
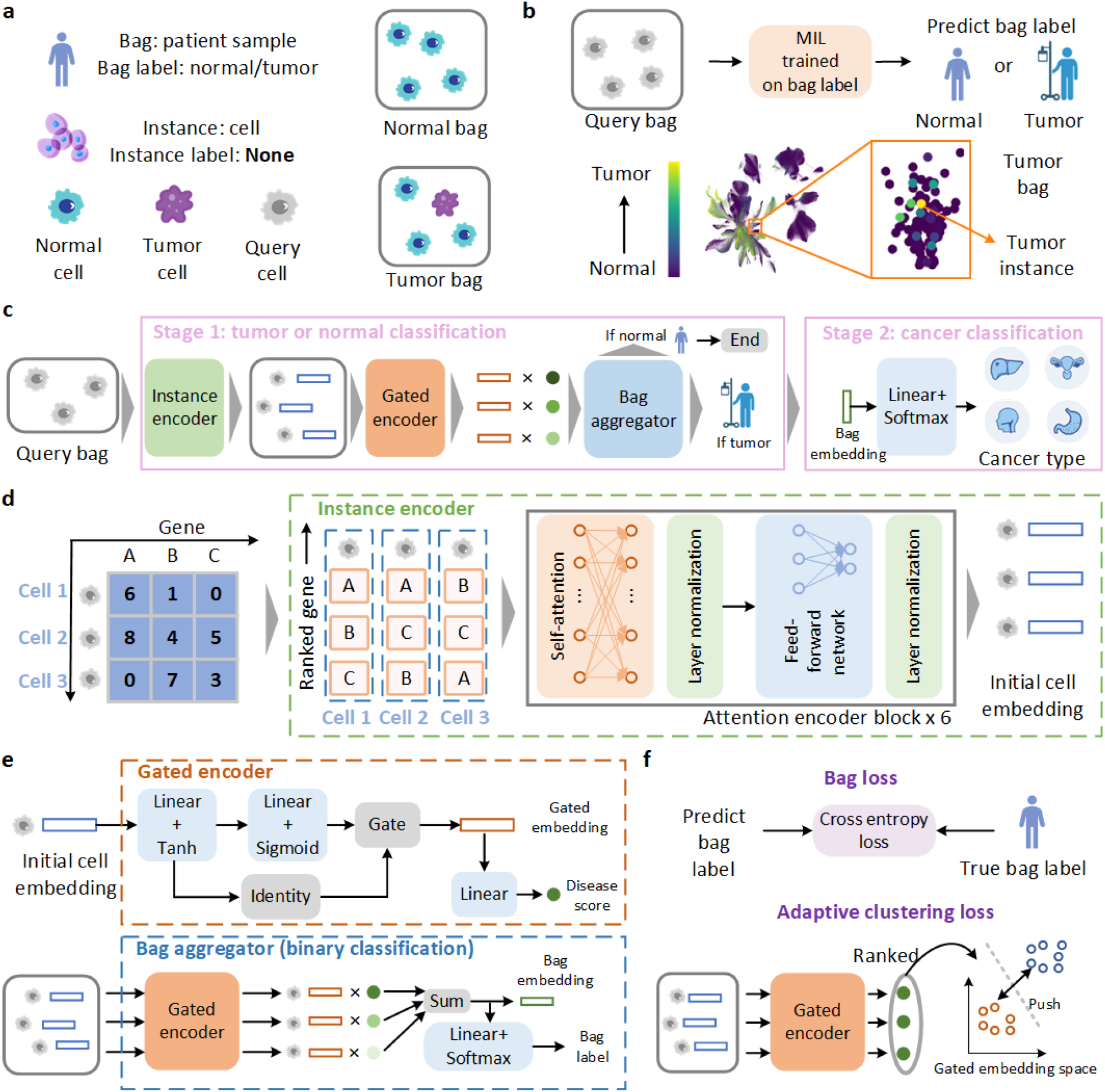
The overall framework of scTransMIL. a. A patient sample (bag) is composed of cells (instances). The cancer phenotype of the patient sample (bag label) is known, while the cancer phenotype of the cells (instance label) is unknown. b. scTransMIL can predict the cancer phenotype of the bag and infer the disease scores of the instances. c. scTransMIL first classifies the patient sample (bag) as tumor or normal, and then further classifies the tumor bag into a specific cancer type. d. Each cell is encoded into a gene sequence. The transformers learn the whole-genome context and output the initial cell representation. e. The initial cell embeddings are integrated into a bag embedding by the gated encoder and the bag aggregator. f. The bag loss and the adaptive clustering loss jointly guide the parameters update to solve the over-fitting problem in the identification of positive instances.

scTransMIL operates in two stages (**Figure 1c**). In *Stage 1*, an *Instance encoder* generates initial embeddings for each cell in the query bag, a *Gated encoder* outputs gated embeddings and instance-level disease scores, and a *Bag aggregator* integrates these gated embeddings of all instances to output a bag embedding, which is classified as tumor or normal. If classified as a tumor bag, it enters *Stage 2*, which employs a *multi-classifier* to predict the specific cancer type using the bag embedding from *Stage 1*.

To ensure accuracy and robustness, scTransMIL incorporates several critical designs. For the *Instance encoder* (**Figure 1d**), the transcriptomic profile of each cell is encoded into a ranked gene sequence, for which batch effect is removed since gene ranking is more robust than expression values. The ranked gene sequence of each cell is fed into a 6-layer Transformer to generate initial cell embeddings. In the *Gated encoder* (**Figure 1e**), a learnable gate filters bag label-irrelevant information from the initial cell embeddings, generating gated embeddings, which are then linearly projected to produce instance-level disease scores. In the *Bag aggregator* (**Figure 1e**), all gated embeddings and disease scores in the bag are weighted and summed into a bag embedding, which is fed into the *classifier* to predict the bag label. During model training, the bag loss and adaptive clustering loss jointly guide the parameters update. The bag loss only requires the bag label and uses cross-entropy to update the entire scTransMIL. Additionally, the adaptive clustering loss sorts gated embeddings by disease scores to maximize separation between the gated embeddings at the extremes of the sorted sequence, thereby mitigating over-fitting in positive instance identification (**Figure 1f**).

### Comprehensive evaluation of tumor prediction across pan-cancer data

We first evaluated the performance of scTransMIL against several baseline methods, including KIDA, DEGAS, scIDST, MultiMIL and two variants of scTransMIL (scTransMIL(PCA) and scTransMIL(FM)) on a pan-cancer dataset^47^. To make fair comparison with MultiMIL, which was originally designed for multi-omic data, we adopted its MIL module for single-omic data analysis by generating input embeddings using scVI. Additionally, we also developed two scTransMIL variants: scTransMIL(PCA) utilizing PCA-reduced cell embeddings and scTransMIL(FM) incorporating cell embeddings derived from a foundation model, both serving as alternatives to our *Instance encoder*’s initial embeddings.

Our evaluation framework employed a comprehensive pan-cancer dataset (comprising 4 million cells and 20,000 patient samples across 30 cancer types, aggregated from 104 public datasets) to rigorously assess the model performance using Accuracy (Acc), Area Under the Receiver Operating Characteristic Curve (AUC), and F1 Score (F1) metrics. The pan-cancer dataset’s remarkable diversity across cancer types provided an ideal testbed for evaluating the model’s generalization capabilities across different tumor profiles. We implemented strict separation of data by randomly splitting the dataset into 80% training, 10% validation, and 10% test sets, ensuring that donors in the test and validation sets were completely unseen during training. This design allows for a rigorous evaluation of scTransMIL’s ability to predict cancer phenotypes from single-cell data, highlighting its robust performance across diverse cancer types and new patient samples.

To evaluate the impact of training data size on model performance, we generated three stratified subsets of different sizes (50%, 75%, and 100%) through random sampling from the aforementioned training set. We trained scTransMIL and the comparison methods on each training subset separately while maintaining a fixed test set for unbiased evaluation. As illustrated in **Figures 2a-c**, all benchmarked methods exhibited performance gains with increasing training data, though scTransMIL demonstrated the most substantial improvements, consistently outperforming the compared methods.

**Figure 2.**
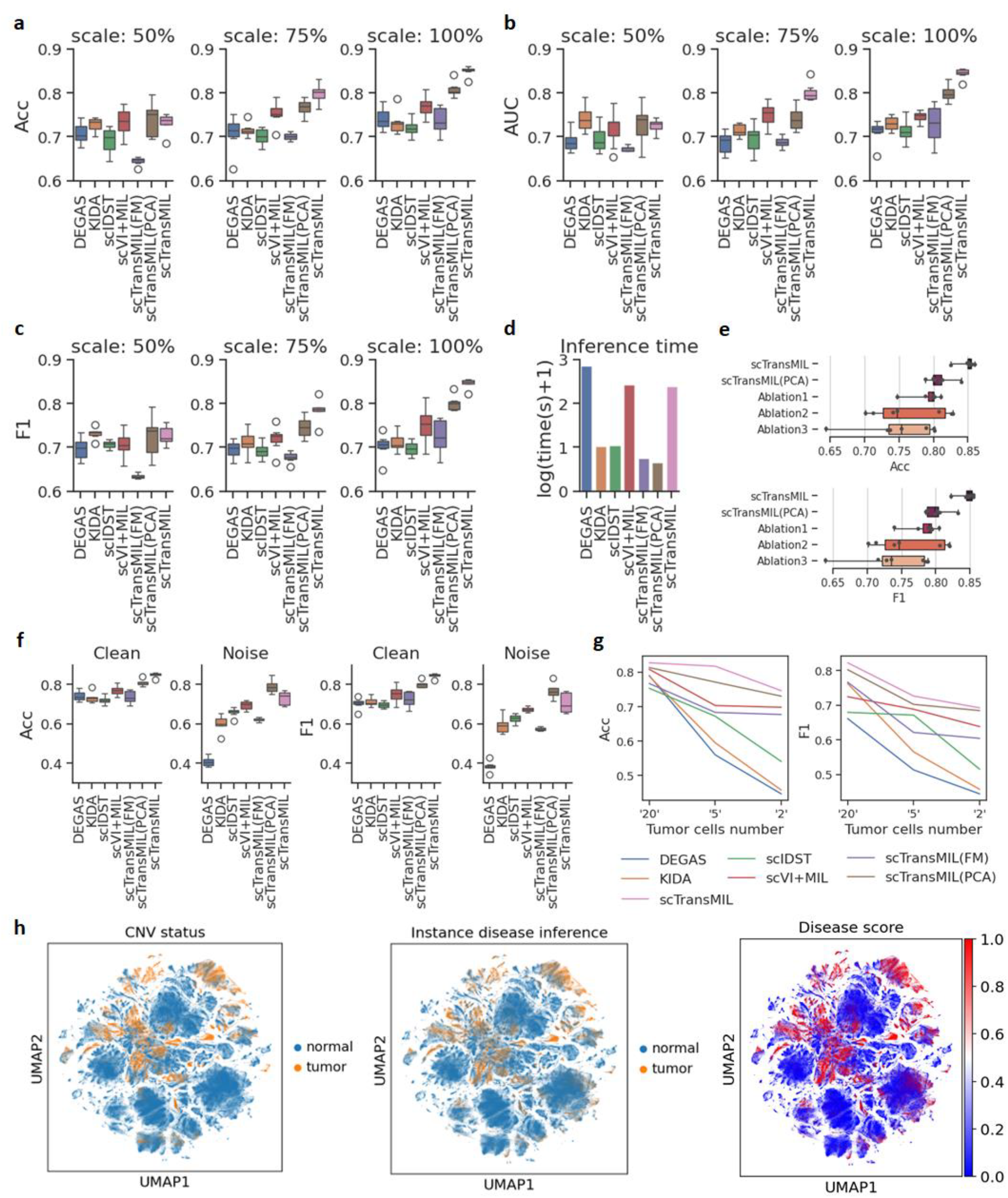
The benchmarks for tumor prediction across pan-cancer data. a. Comparison of Acc (Accuracy) under different training scales. b. Comparison of AUC (Areas Under the Curve) under different training scales. c. Comparison of F1 (F1 Scores) under different training scales. d. Comparison of the average inference time on a single bag. e. Ablation experiments, with metrics including Acc and F1. Ablation1: removing the adaptive clustering loss. Ablation2: removing the gating mechanism. Ablation3: replacing the attention pooling with max pooling. f. Comparison between the noisy setting (Noise) and the default setting (Clean), with metrics including Acc and F1. There are n = 7 repeats with different random seeds for data splitting. Noisy setting: if a donor was diagnosed with cancer, then all sample bags from this donor were assigned tumor labels, regardless of the actual composition of individual samples. g. Comparison of prediction results in the simulation scenario of early screening, with the metrics including Acc and F1. The number of tumor cells in the samples decreases successively from 20 to 5 and then to 2. The fewer the tumor cells are, the greater the difficulty of early cancer screening will be. h. Inferred disease-relatedness at the single-cell resolution. From left to right are the reference atlas (CNV status), the single-cell cancer phenotypes inferred by the model (Instance disease inference), and the disease scores inferred by the model (Disease score).

To evaluate computational efficiency in real-world large-scale applications, we benchmarked the inference time required for processing a single bag using scTransMIL and the compared methods. As shown in **Figure 2d**, both DEGAS and scVI+MIL showed prolonged processing time, primarily attributable to their computationally intensive data integration pipelines. While scTransMIL similarly exhibited extended inference time due to its whole-genome transcriptome processing and embedding generation requirements, the scTransMIL(PCA) variant demonstrated markedly improved efficiency. By leveraging PCA-reduced embeddings in place of the full instance encoder, scTransMIL(PCA) significantly reduced inference time while maintaining competitive performance. This positions scTransMIL(PCA) as a particularly advantageous and practical choice for large-scale deployment and application scenarios requiring high-throughput processing.

To further validate the importance and effectiveness of scTransMIL’s key design elements, we conducted ablation experiments through removing the adaptive clustering loss (Ablation1), gating mechanism (Ablation2), or replacing the attention pooling with max pooling (Ablation3) (**Extended Data Figure 1**). As shown in **Figure 2e**, removal of any of these key components resulted in a decline in model performance. Notably, models using whole-genome data as input consistently outperformed those relying on PCA-reduced embeddings, highlighting the value of preserving whole-genome information for cancer phenotype inference.

Additionally, to rigorously assess model robustness under challenging real-world conditions, we introduced noisy bag labels by assigning labels at the donor level—a scenario reflecting clinical settings where sample-level annotations may be unavailable. Under this setting, all samples from a cancer-diagnosed donor were uniformly labeled as tumor-positive, regardless of their actual cellular composition, thereby artificially introducing noise into data labels. Comparative analysis between this noisy setting (Noise) and the default setting (Clean) (**Figure 2f**) revealed that MIL-based approaches (scVI+MIL, scTransMIL and its variants) maintained remarkable stability and resistance to label noise across all key metrics. These results provide empirical validation that MIL frameworks are particularly well-suited for bridging patient-level cancer phenotypes with single-cell data, as they can maintain high fidelity predictions even when the label noise commonly encountered in clinical datasets.

Moreover, to evaluate scTransMIL’s potential for early cancer detection, we performed simulation experiments under varying degrees of tumor cell scarcity, gradually decreasing the number of tumor cells per sample from 20 to 5 and finally to just 2 cells (**Figure 2g**). scTransMIL consistently maintained superior performance compared to all compared methods across both Acc and F1 score, demonstrating robust predictive capability even in scenarios with extremely limited malignant cells. Notably, while all methods experienced performance decline as tumor cell counts decreased, scTransMIL exhibited the least degradation, underscoring its effectiveness and reliability for early cancer detection, especially in challenging clinical contexts with minimal tumor presence.

To rigorously evaluate scTransMIL’s ability to infer disease-relatedness at the single-cell resolution, we performed comprehensive validation of its disease-relatedness predictions across the entire 4 million-cell pan-cancer atlas. As shown in **Figure 2h**, we visualized single-cell disease scores predicted by scTransMIL by setting the threshold of 0.5 to assign tumor or normal labels to individual cells (Instance disease inference). We observed striking concordance between scTransMIL’s predictions and gold-standard CNV-derived annotations (CNV status). The results indicate that scTransMIL achieved highly accurate cancer phenotype inference across the 4 million single-cell pan-cancer atlas, establishing it as a reliable framework for single-cell phenotypic inference at scale.

### Prediction of cancer type and generalization to out-of-distribution data

Building upon its accurate tumor/normal classification, scTransMIL’s *multi-classifier* module demonstrates exceptional capability in determining specific cancer types—a feature with particular clinical relevance and value for challenging clinical applications like Cancer of Unknown Primary (CUP). CUP presents a significant diagnostic dilemma, where metastatic lesions are identified without detectable primary tumors. This diagnostic uncertainty poses significant clinical challenges, as accurately identifying the tumor origin is critical for determining optimal treatment strategies^48^ yet remains particularly difficult in such cases. Leveraging the comprehensive cancer type annotations provided in our pan-cancer dataset (30 distinct types; **Extended Data Figure 2a**), scTransMIL achieved remarkable classification accuracy for both primary and metastatic tumors in held-out test samples (**Figures 3a-b**). The model’s robust performance on metastatic specimens is especially noteworthy, as it directly addresses a critical unmet need in clinical diagnosis and management of CUP.

**Figure 3.**
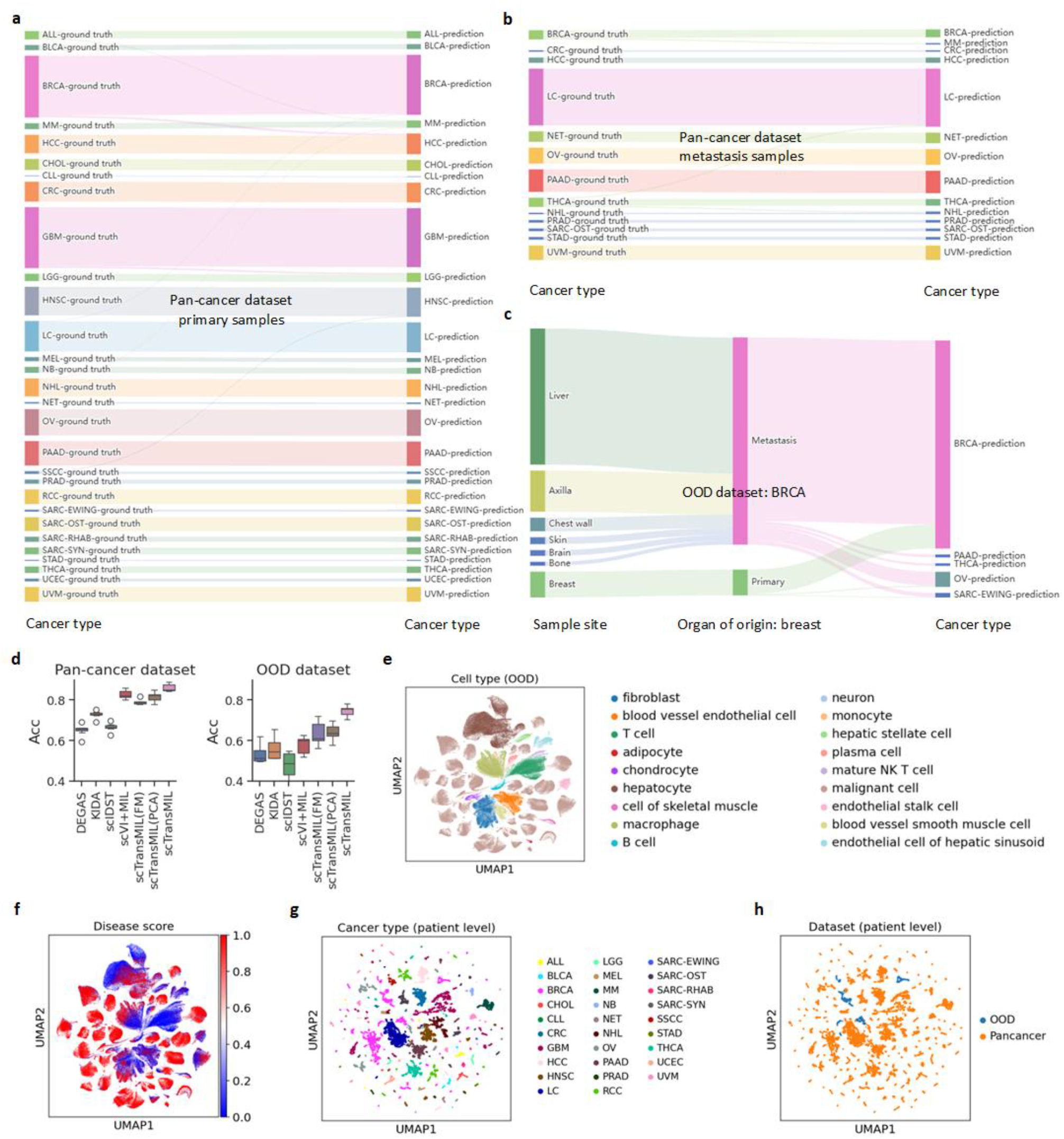
The evaluation of cancer type prediction and generalization to OOD data. a. The cancer type prediction results for the subset of primary samples in the pan-cancer dataset. b. The cancer type prediction results for the subset of metastatic samples in the pan-cancer dataset. c. The cancer type prediction results for the OOD breast cancer dataset. d. The prediction performances of different methods on the pan-cancer dataset and the OOD breast cancer dataset. There are n = 7 repeats with different random seeds for data splitting. e. The cell-level atlas of the OOD breast cancer dataset, colored according to the cell types. f. The cell-level disease scores inferred by the model for the OOD breast cancer dataset. g. The bag embedding space output by scTransMIL on the pan-cancer dataset and the OOD dataset, colored according to the cancer category. The bags in the pan-cancer dataset and the OOD dataset were jointly embedded into a common space. h. The bag embedding space output by scTransMIL on the pan-cancer dataset and the OOD dataset, colored according to the data source.

To rigorously evaluate scTransMIL’s generalizability, we tested its performance on an independent Out-Of-Distribution (OOD) breast cancer (BRCA) dataset^49^ comprising 400,000 cells from 1,500 tissue samples across 37 patients (**Extended Data Figure 2b**). In this dataset, all samples were malignant breast cancer and accordingly all bags were assigned the BRCA label. As a result, scTransMIL demonstrated robust classification accuracy with almost all primary samples correctly identified as BRCA, and with only a small fraction of metastatic samples misclassified as other cancer types (**Figure 3c**), presumably due to the complexity of OOD data and metastatic samples.

Our comparative analysis revealed scTransMIL’s superior generalization capability through benchmarking experiments against state-of-the-art methods on both the pan-caner test set and the OOD breast cancer dataset (**Figure 3d**). While all the comparison methods showed significant performance degradation when applied to the OOD dataset, scTransMIL maintained accurate and robust classification performance, demonstrating excellent generalization capability. Beyond cancer type prediction, scTransMIL could also infer the disease scores at the single-cell resolution to the OOD data. As shown in **Figures 3e-f**, cells with high disease scores assigned by scTransMIL showed high concordance with malignant cells in OOD data, demonstrating robust cross-dataset generalization at single-cell resolution.

To elucidate scTransMIL’s remarkable generalization capability, we conducted a comprehensive latent space representation analysis across the pan-cancer dataset and the OOD dataset, by extracting and visualizing all the bag-level embeddings generated by the model’s *Bag aggregator*. The UMAP^50^ visualizations revealed three key findings: (1) The learned representations formed distinct, cancer-type-specific clusters (e.g., distinct clusters between lung cancer (LC) and colorectal cancer (CRC) populations) in both pan-cancer and OOD datasets (**Figure 3g**), demonstrating preservation of biologically meaningful patterns; (2) The samples in the OOD dataset showed seamless integration with corresponding cancer clusters from the pan-cancer dataset (**Figure 3h**), indicating that scTransMIL’s exceptional ability to effectively mitigate batch effects and dataset-specific biases. Altogether, these findings collectively demonstrate that scTransMIL learns robust, biologically informed representations that not only enable accurate cross-dataset generalization but also possess clinical utility for patient stratification and precision oncology applications.

### Identification of tumor-specific genes in ALL using scTransMIL

scTransMIL’s attention mechanism leverages the rich whole-genome context learned from the pan-cancer dataset to provide comprehensive gene-level biological insights, enabling systematic identification of cell type-specific genes across the complete genome. Our analysis of acute lymphoblastic leukemia (ALL) patient samples, which form distinct clusters in the embedding space of the pan-cancer dataset (**Figure 4a**), revealed remarkable concordance between scTransMIL’s single-cell disease scores and orthogonal corrected CNV-based malignancy annotations (**Figures 4b-c**), with malignant cells predominantly mapping to the B cell lineage (**Figure 4d**). Focusing on matched tumor (all_GSE132509_ETV6_RUNX1_4) and normal (all_GSE132509_HHD_2) B-cell populations (**Extended Data Figure 3**), scTransMIL’s attention-score ranking not only uncovered the established ALL markers like CD79A and PIM1^51^ within the top 100 genes but also uniquely identified BLK, a known tumor B-cell marker that conventional methods miss (**Figures 4e-f**).

**Figure 4.**
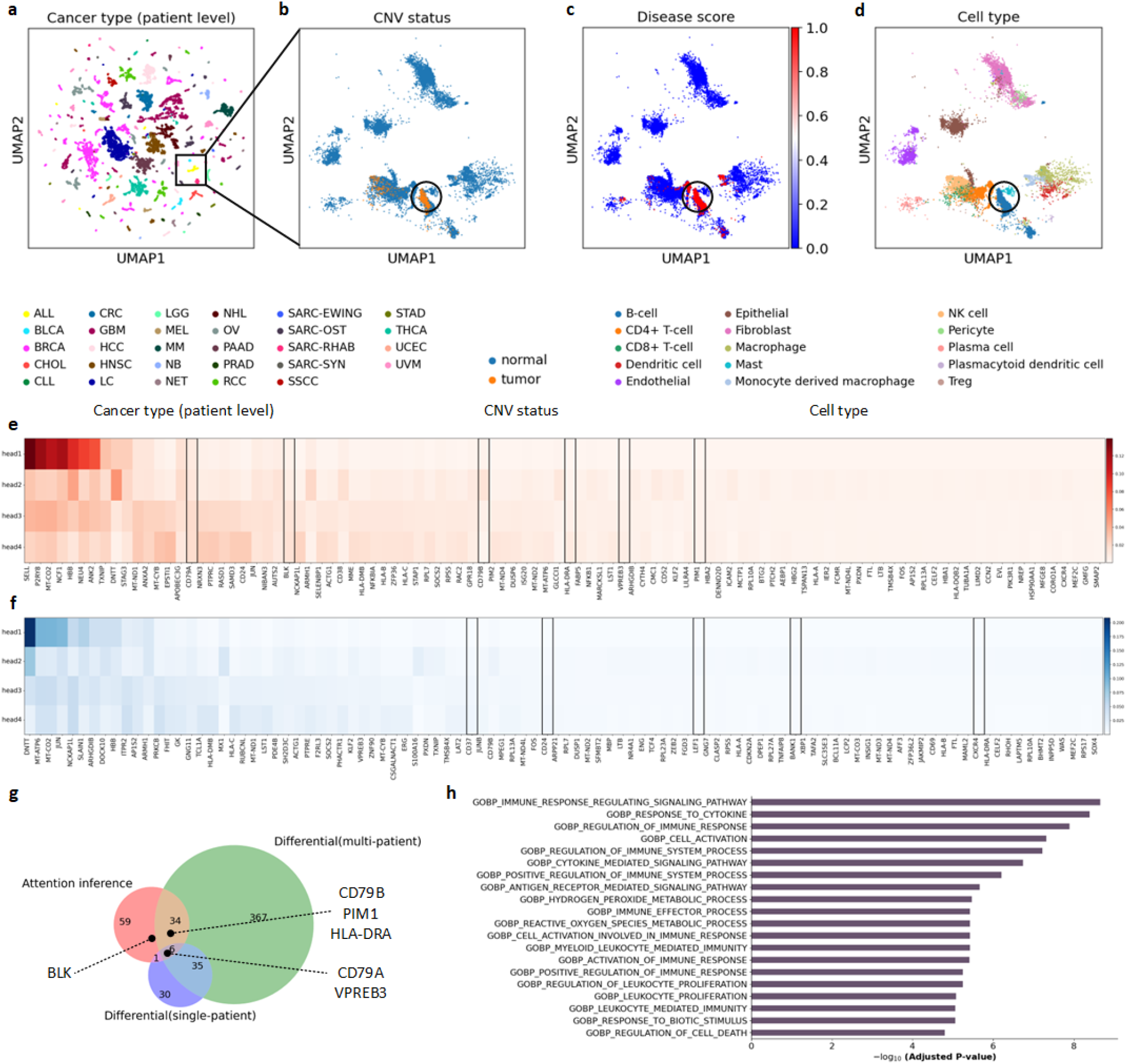
Tumor-specific gene inference by scTransMIL. a. In the bag embedding space, the black box represents a selected batch of ALL bags. b. The single-cell atlas corresponding to the selected ALL bags, colored by CNV annotation. c. The single-cell atlas corresponding to the selected ALL bags, colored by disease scores. d. The single-cell atlas corresponding to the selected ALL bags, colored by cell types. e. The top 100 genes specific to tumor B cells inferred by scTransMIL. The black box indicates the specific genes of ALL B cells known from previous studies. The rows of the heatmap represent the 4 heads, the columns represent the gene names. f. The top 100 genes specific to normal B cells inferred by scTransMIL. The black box indicates the specific genes of normal B cells that are already known. The rows of the heatmap represent the 4 heads, the columns represent the gene names. g. Comparison of the gene sets inferred by scTransMIL attention mechanism and differential analysis. Six known specific genes of ALL B cells (reference genes) are marked with text. h. The enrichment analysis of the top 100 genes specific to tumor B cells inferred by scTransMIL, with the top 20 pathways visualized.

Our systematic comparison between scTransMIL’s attention-based inference and traditional differential expression analysis (DEA) methods reveals several key advantages. When analyzing B cells from a single ALL patient (all_GSE132509_ETV6_RUNX1_4), conventional DEA identified only CD79A and VPREB3 due to inherent sample size limitation and sample selection bias, while multi-patient DEA (incorporating B cells from all ALL patients in the pan-cancer dataset) still failed to detect BLK—a well-established tumor B cell marker—despite increased statistical power (**Figure 4g**). scTransMIL uniquely recovered BLK through its whole-genome attention mechanism, even when analyzing individual samples. This capability stems from scTransMIL’s capacity to recognize biologically significant genes through their contextualized genomic patterns rather than relying solely on differential expression, as confirmed by BLK’s constant expression levels between tumor and normal samples (**Extended Data Figure 4)**. The model’s ability to identify such functionally important yet non-differential genes demonstrates its superiority over conventional approaches for comprehensive biomarker discovery.

To biologically validate the functional relevance of the identified gene sets by scTransMIL, we performed comprehensive pathway enrichment analysis, which revealed that the top 20 enriched pathways (FDR<0.001) were strongly associated with ALL pathogenesis (**Figure 4h, Supplementary Table 1**), including immune response regulating signaling pathway, cytokine-mediated signaling pathway, and antigen receptor-mediated signaling pathway. The robust enrichment of these biologically coherent pathways provides strong validation that scTransMIL’s attention mechanism captures functionally relevant gene sets beyond what conventional differential expression analysis can identify, confirming its ability to extract meaningful biological insights from single-cell transcriptomic data.

### Deciphering novel AML heterogeneity and subtypes with scTransMIL

To evaluate scTransMIL’s generalization capability on novel diseases beyond its original 30-class cancer types, we performed systematic fine-tuning experiments on an acute myeloid leukemia (AML) dataset— a disease category absent from the pan-cancer training data. While scTransMIL is pre-trained to recognize 30 cancer types, its ability to adapt to previously unseen diseases is essential for expanding its applicability. The AML dataset comprised 35 tumor and 6 normal samples (**Extended Data Figure 5a**), with gold-standard cell-level malignant annotations generated through an improved seq-well measurement technique in the original study, which is of high-cost and inefficient^52^. In contrast to these costly single-cell labelling methods, scTransMIL demonstrated remarkable data efficiency by achieving robust performance using only a few bag-level tumor/normal labels for fine-tuning. We implemented a strategic binary classification framework during fine-tuning to prioritize fundamental tumor-normal discrimination while preserving the model’s capacity to uncover novel biological heterogeneity^53^. To identify the most effective fine-tuning strategy, we compared three experimental settings: i) direct application of the pre-trained model without fine-tuning; ii) fine-tuning using only the *Bag aggregator* while freezing the whole *Instance encoder*; and iii) fine-tuning the last two transformer blocks and the *Bag aggregator*, while freezing the remaining layers. As shown in **Figure 5a**, we found that selectively fine-tuning the last two transformer blocks alongside the *Bag aggregator* (while freezing fewer parameters) yielded optimal performance on the novel diseases.

**Figure 5.**
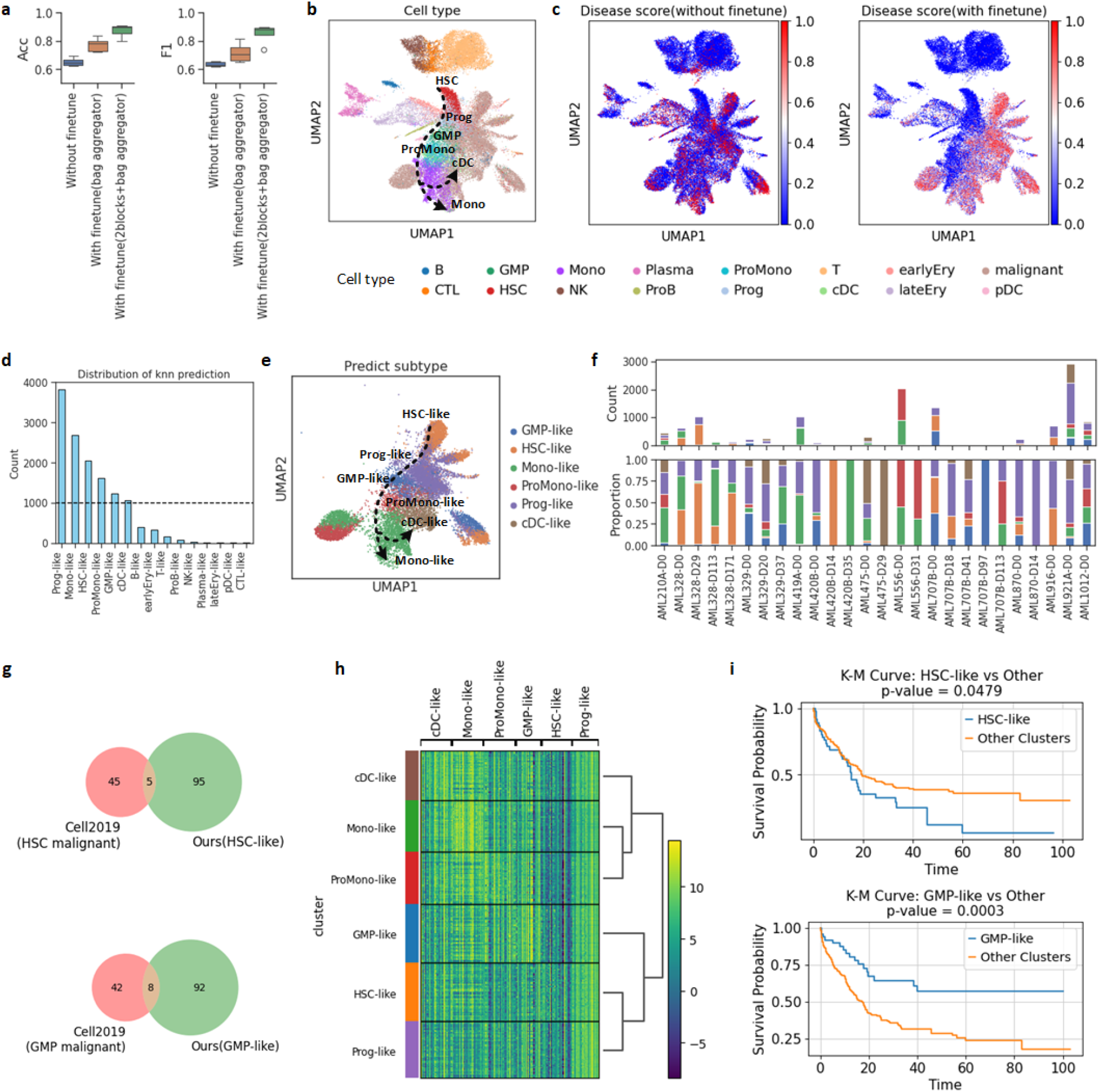
scTransMIL enables novel cancer heterogeneity analysis on AML. a. Prediction performance of AML tumor bags before and after fine-tuning scTransMIL. b. Visualization of the single-cell atlas based on instance embeddings generated by scTransMIL, colored by cell types. The arrows indicate the differentiation trajectory of normal hematopoietic cells. c. Disease scores corresponding to the model before or after fine-tuning. d. Cell number statistics of AML subtypes inferred by scTransMIL. e. Single-cell atlas (tumor cells only) of AML subtypes inferred by scTransMIL. f. Total cell number (top) and proportions (bottom) of AML subtypes in different patients at different treatment stages. g. Comparison between the subtype-associated genes recommended by scTransMIL and those recommended by the original study, conducted on HSC-like (top) and GMP-like (bottom) subtypes respectively. h. Validation of the AML subtype-specific genes recommended by scTransMIL using bulk AML RNA-seq data. The heatmap shows the differential expression of bulk samples grouped based on specific genes. The horizontal axis represents the subtype-specific genes inferred by scTransMIL, and the vertical axis represents the subtype samples redefined by subtype-specific genes. i. Survival analysis using the death time provided by the bulk data, comparing HSC-like (top) and GMP-like (bottom) subtypes respectively.

To interpret the representation ability of scTransMIL, we extracted the instance embeddings of all cells from the AML dataset and compared with the original single-cell expression profiles. Notably, the fine-tuned scTransMIL model generated embeddings that captured a continuous hematopoietic differentiation trajectory from hematopoietic stem cells (HSC) to monocytes (Mono) in normal bone marrow samples (**Figure 5b**)—a biologically meaningful pattern not discernible in the original single-cell expression profiles (**Extended Data Figure 5b**). Furthermore, disease scores inferred by the post-fine-tuned scTransMIL model showed markedly improved alignment with the ground truth malignancy annotations compared to the pre-fined-tuned model (**Figure 5c**), demonstrating scTransMIL’s ability to accurately differentiate tumor and normal cells using only sample-level labels. Expanding beyond binary classification, we identified six distinct AML subtypes (each comprising >1000 tumor cells) (**Figure 5d**), by applying a conservative disease score threshold of > 0.2 followed by kNN-based annotation relative to normal differentiation states. These six AML subtypes identified by scTransMIL precisely recapitulated the established AML differentiation trajectory, faithfully capturing the continuum from primitive malignant HSCs through GMP-like to terminally differentiated Mono-like cells (**Figure 5e**). This recapitulation of known biological trajectories^52^ provides strong validation of scTransMIL’s ability to resolve clinically relevant heterogeneity. Longitudinal analysis of AML patient samples across treatment timepoints revealed clinically meaningful shifts in tumor cell burden and subtype composition during treatment, exemplified by two representative cases: patient AML707B maintained predominantly GMP-like subtype, consistent with low myeloid differentiation marker expression indicated by flow cytometry in the original study^52^, while patient AML419A showed Mono-like subtype, aligning with high myeloid differentiation marker expression (**Figure 5f**). These findings collectively demonstrate scTransMIL’s unique capability to simultaneously resolve fundamental biological processes (normal differentiation), disease states (malignant transformation), clinically relevant tumor evolution trajectory and relevant shifts in patient treatment response.

Moreover, to validate the clinical relevance of scTransMIL’s AML subtyping, we performed a comprehensive survival analysis on an independent cohort of 671 AML patients^54^ containing bulk RNA-seq data only. Subtype-specific gene sets (top 100 genes per subtype) identified by the attention mechanism of scTransMIL (**Extended Data Figure 6**) showed partial overlap with those established marker genes in the original single-cell AML study^52^ while revealing novel subtype-associated genes (**Figure 5g**). When applied to bulk AML RNA-seq data, these gene signatures enabled clear stratification of patients into distinct prognostic groups through hierarchical clustering (**Figure 5h**), with primitive (HSC-like) and differentiated (GMP-like/Mono-like) subtypes forming separate branches. Kaplan-Meier survival analysis was conducted on each subtype of AML patients, and showed that HSC-like patients exhibited significantly worse prognosis compared to GMP-like cases (**Figure 5i, Extended Data Figure 7**). This finding is consistent with known clinical knowledge that malignant HSCs are more resistant to chemotherapy, evade immune clearance, and contribute to relapse^55^. Functional enrichment analysis of subtype-specific genes provided mechanistic explanations for these clinical observations: HSC-like AML-specific genes were strongly associated with immune evasion pathways and correlated with poor survival, e.g., “GSE40666 UNTREATED VS IFNA STIM STAT1 KO CD8 TCELL 90MIN DN” (**Supplementary Table 2, Extended Data Figure 8**), while GMP-like AML-specific genes were associated with differentiation pathways and correlated with better treatment response. These results collectively illustrate scTransMIL’s unique ability to provide biologically meaningful, mechanistic and clinically actionable insights into novel uncharacterized cancers, tumor heterogeneity, treatment response dynamics, and interpretable prognostic markers from single-cell data, establishing it as a promising framework for data-driven translational cancer research that bridges single-cell resolution with clinical outcomes.

## Discussion

The high resolution of single-cell sequencing data can bring about significant changes to the clinical diagnosis of cancer and the research on its mechanisms. Currently, patient-level phenotypic labels are the available meta-information in the samples. However, methods based on bulk data cannot detect malignant cells at the single-cell resolution, making it difficult to apply them in the early screening of cancer and the inference of its mechanisms. At the same time, accurate phenotypic annotation at the single-cell level is still lacking, which also hinders the application of existing extensive single-cell data-based methods in the early screening of cancer and the analysis of its heterogeneity. In order to connect patient-level phenotypes with single-cell transcriptome data, we proposed scTransMIL for single-cell data based on multi-instance learning.

The scTransMIL proposed in this study is a deep learning model based on single-cell transformer and multi-instance learning, which can effectively connect the cancer phenotypes of patient samples with single-cell atlases. scTransMIL achieves 3 functions simultaneously: 1) Given a sample from a donor, accurately predict the cancer phenotype of this sample for early diagnosis. 2) Infer the disease-relatedness of each cell in the sample at the single-cell resolution to study the cellular processes driving the disease phenotype. 3) Infer genes specific to tumor cells across the whole genome to promote research on potential biomarkers.

Extensive experiments proved the versatile applicability and superior performance of our method. scTransMIL has achieved state-of-the-art performance in benchmark tests. scTransMIL can make correct predictions in samples containing only one or two malignant cells. This result breaks through the limitations of bulk data in the current clinical practice, and it can provide assistance for the early screening of cancer. scTransMIL can accurately predict the cancer types in out-of-distribution datasets and performs excellently even in predicting metastatic samples, which is of great significance for the clinical diagnosis of cancers of unknown primary origin. In addition, the disease-relatedness inferred by scTransMIL at the single-cell resolution is highly consistent with the ground truth of malignant cells. In terms of biological insights, scTransMIL has significant advantages. Traditional gene-level analysis methods rely on differential analysis and are easily affected by sample selection, while scTransMIL takes into account the whole-genome space, and its attention mechanism can provide global gene-level biological insights. In the tumor-normal analysis of samples from patients with acute lymphoblastic leukemia, scTransMIL successfully recommended markers for tumor B cells and normal B cells. For the analysis of new cancer types, scTransMIL shows good adaptability and interpretability. Taking acute myeloid leukemia as an example, minimal fine-tuning scTransMIL using only a few patient sample-level labels enabled: 1) reconstruction of differentiation trajectories, 2) identification of clinically distinct AML subtypes at the single-cell resolution. By integrating these capabilities—precise single-cell classification, interpretable gene-level insights, and adaptation to novel diseases—scTransMIL will emerge as a powerful and transformative tool for precision oncology, with direct applications in early cancer detection, tumor heterogeneity dissection, and personalized treatment strategy design.

## Methods

In this section, we first introduce the model architecture and loss function of scTransMIL. Then, we present the usage of scTransMIL, including training, inference, and fine-tuning. Next, we introduce the datasets and comparative methods involved in the experiments. Finally, we provide the detailed information of the complete application setup. scTransMIL first classifies patient samples into tumor bags or normal bags (the first stage) and further distinguishes the cancer type of tumor bags (the second stage). In the first stage, the instance encoder generates the initial embedding for each instance, the gated encoder outputs the gated embedding and disease score for each instance, and the bag aggregator integrates the gated embeddings to obtain the bag embedding for binary classification. For tumor bags, the second stage employs a multi-classifier to predict the cancer type. During training, the model is optimized using both bag loss and adaptive clustering loss.

### Instance encoder

Given a patient sample (bag), which contains multiple single cells, the instance encoder generates an initial embedding for each single cell. For a single cell, we use rank value encoding to convert the gene expression levels of the cell into a gene sequence. The gene sequence can be regarded as a sentence, with genes acting as words. Rank value encoding takes into account the whole-genome characteristics of the single cell. We filter out genes with an expression level of 0, thus adaptively reducing the length of the gene sequence for each single cell. Then, we rank the remaining genes according to their gene expression levels to obtain a gene sequence (genes, as words, are mapped to IDs in the vocabulary). Since the gene sequence does not involve specific expression levels, batch effects can be alleviated. The instance encoder consists of 6 transformer encoder units, and each unit is composed of a self-attention layer and a feed-forward neural network layer^56^. The instance encoder transforms genes into a 256-dimensional space through the transformer. This space encodes the characteristics of the gene in a specific cellular environment (gene sequence) through self-attention. The initial embedding of any cell is generated by averaging the embeddings of each gene in the gene sequence, resulting in a 256-dimensional initial cell embedding. For scTransMIL(PCA) and scTransMIL(FM), we use the PCA cell embedding and the cell embedding output by the foundation model respectively to replace the initial cell embedding output by the instance encoder.

### Gated encoder

In a bag, based on the initial cell embeddings, the gated encoder updates the cell embeddings through a gating mechanism, and outputs the gated embeddings and disease scores. The gating mechanism can filter out the noise signals in the initial cell embeddings that are irrelevant to the category of the bag^57^.

Let the initial cell embedding of cell (instance) *i* be 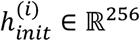, and the corresponding gated embedding and disease score be 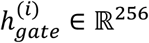 and *a*^(*i*)^ ∈ ℝ respectively.

The initial cell embedding of instance *i* is input into the gated encoder. First, it passes through a non-linear transformation module composed of a linear transformation layer and a tanh activation function to obtain the updated cell embedding 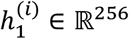.Then, it goes through another non-linear transformation module consisting of a linear transformation layer and a sigmoid activation function to output the filtering vector 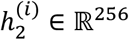.The filtering vector acts on the updated cell embedding and outputs the gated embedding, which is used to retain informative features and suppress non-informative features:

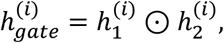

where ⊙ represents element-wise multiplication. Based on the binary classification scenario in the first stage, each bag will be assigned either a tumor label (1) or a normal label (0). We use the disease score to measure the correlation between cell *i* and the tumor label. We input the gated embedding into *linear*(*·*) and output the disease score:

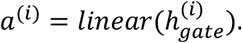

### Bag aggregator

For a bag *j*, assume that it contains *m* instances (the number of instances in different bags is variable). We use the instance encoder and the gated encoder to obtain the gated encoder to obtain the gated embeddings 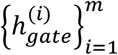 and disease scores 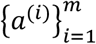 of all instances in the bag. The bag aggregator is used to integrate the instance representations in the current bag *j* and output a bag embedding 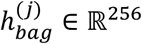.We achieve the integration by weighting the gated embeddings with the disease scores:

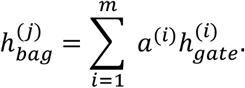

In the first stage, the bag embedding is input into a linear transformation and a binary softmax activation function to obtain the binary classification result 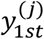.The bag will be assigned either a tumor label or a normal label. If the bag is assigned the tumor label, the bag embedding is input into another linear transformation and a multi-class softmax activation function to obtain the multi-classification result 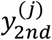, and the bag will be assigned a cancer type.

### Loss

The model is trained under the assumption of multiple-instance learning (MIL), which is a weakly supervised learning process. We only need to provide the bag labels. With the goal of bag classification, the model can infer the correlation between instances and bag labels by updating its parameters. For bag-level supervision, let the true label of bag *j* be 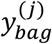.Taking the binary classification of tumor or normal as an example, we define the bag-level loss 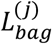 using the binary cross-entropy loss:

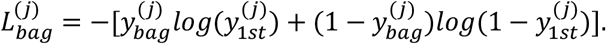

For the cancer type identification in the second stage, we use the multi-class cross-entropy loss:

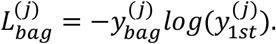

To prevent the model from overfitting on a few easily classifiable positive instances, the adaptive clustering loss is used to assist the model in distinguishing between positive and negative instances. To balance the learning of positive and negative embeddings, we construct the adaptive clustering loss based on the SVM loss. Since the true instance-level labels do not exist, we assign pseudo-labels according to the disease scores. A high disease score indicates a high probability that the instance is positive, while a low disease score indicates a high probability that the instance is negative^41^.

For bag *j*, its true binary classification label is 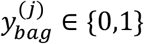,where 1 represents tumor and 0 represents normal. All the disease scores in this bag are sorted. We select the *B* instances with the lowest disease scores and assign them negative labels, and select the *B* instances with the highest disease scores and assign them positive labels.

For an instance *i* in bag *j*, we add a linear transformation *W*_*inst*_ ∈ ℝ^2×256^ to define the instance’s clustering assignment probability as *P*^(*i*)^:

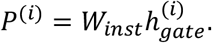

We adopt the smoothed SVM loss. Given that the predicted probability of instance *i* in bag *j* is 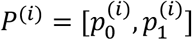 and the assigned pseudo-label is 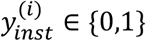,the SVM loss is:

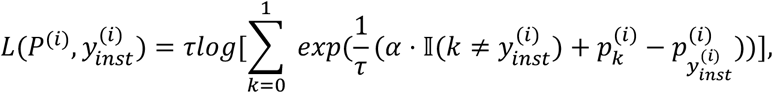

where *τ* is the temperature coefficient, with a default value of 1.0, which is used to control the smoothness of the SVM loss function to facilitate gradient updates. *α* is the specified margin, with a default value of 1.0, which is used to control the interval between the predicted scores of different classes. *II* is the indicator function, which takes the value of 1 when 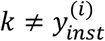,and 0 otherwise.

Since instances with high disease scores do not necessarily all correspond to malignant cells, and instances with low disease scores do not necessarily all correspond to benign cells, there must be noise in the labels created for supervising the instance clustering task during the training process. However, the SVM loss can reduce overfitting when the data labels are noisy.

For any bag *j*, we only need to select 2*B* instances to calculate the adaptive clustering loss 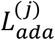:

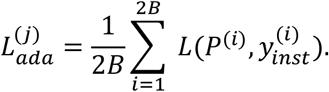

Therefore, the loss on bag *j* is:

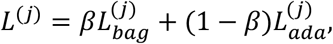

where *β* is the coefficient of the loss function, with a default value of 0.7. We need to prioritize the classification of patient samples.

### Training

We trained scTransMIL on a pan-cancer dataset composed of 104 publicly available cancer datasets^47^. The training resources covered 20,000 patient samples, 4 million single cells, and 30,000 genes. Only the patient sample level (tissue sample level) had cancer phenotype labels, and the cancer phenotype labels included binary classification scenario (tumor or normal) and multi-classification scenario (cancer type). We regarded tissue samples (patient samples) as bags. Each donor usually had multiple tissue samples. Therefore, there were two ways to assign labels to bags. For the first label assignment method, each bag used its corresponding label, and the labels in this way were completely clean. For the second label assignment method, we assigned labels at the donor level. If a donor was a cancer patient, all bags belonging to this donor were assigned the tumor label. This method would introduce noise and could be used to test the model’s learning ability in extreme situations, such as whether the model focused on causal features instead of statistically correlated features. scTransMIL was trained and inferred in the first label assignment method by default.

The dataset was randomly divided into a training set (80% of the donors), a validation set (10% of the donors), and a test set (10% of the donors). This ensures that the donors in the test set or validation set do not overlap with those in the training set, thus directly evaluating the generalization ability of the model on unseen donors.

During training, bags were randomly sampled with a batch size of 1. The multi-nomial sampling probability of each bag was inversely proportional to the frequency of its true category to mitigate the class imbalance problem in the training set. The model parameters were randomly initialized and trained end-to-end using bag-level labels. The model parameters were updated by the Adam optimizer with a learning rate of 2 × 10^−4^ and a weight decay of 1 × 10^−5^. The model was trained for at least 30 epochs. If the early stopping criterion was not met, it was trained for a maximum of 200 epochs. We monitored the validation loss during training. If the validation loss did not decrease for 10 consecutive epochs, we used early stopping. We selected the model with the lowest validation loss for testing on the test set.

### Inference

Given a bag, scTransMIL can predict whether the bag is a tumor sample or a normal sample and infer the disease score of all instances in the bag. If a bag is predicted to be a tumor sample, scTransMIL further predicts the cancer type of the bag. We first tested the binary classification of scTransMIL on pan-cancer dataset (internal dataset), and then selected all bags with true tumor labels to test the performance of scTransMIL in predicting cancer types in the second stage. To further test the generalization ability of the model on out-of-distribution dataset (OOD dataset), we prepared a new dataset as the test set. This dataset contains 1,500 tissue samples from breast cancer patients^49^.

In addition, we utilized the attention mechanism of scTransMIL to infer genes specific to the target cell group. scTransMIL returns the attention of the last transformer layer corresponding to each instance from the instance encoder, and we calculate the mean of the attention in all heads. After collecting all target instances, we averaged the mean attention of all target instances as the attention specific to the target cell group. According to the attention values, we ranked the genes and selected the top 100 genes as the genes specific to the target cell group.

### Fine-tuning

Given that the pan-cancer training set encompasses 30 types of cancers, for unseen cancer type such as acute myeloid leukemia (AML), we can perform analysis of new disease by fine-tuning scTransMIL. Taking the AML dataset as an example, this dataset includes 5 healthy donors and 16 AML patients^52^. The annotation information used for fine-tuning only comes from the cancer phenotypes (tumor or normal) of these 21 donors. We let scTransMIL only perform fine-tuning at the binary classification stage to avoid overfitting. We froze the parameters of the instance encoder, set the maximum number of training epochs to 30, and kept other settings consistent with those in the pan-cancer training.

### Data

#### Pan-cancer dataset

In the pan-cancer dataset^47^, 20,000 patient samples were collected, including single-cell transcriptome sequencing data of 4 million cells. The feature space covers 30,000 genes. The dataset is composed of 104 publicly available datasets and encompasses 30 different cancer types. Each patient sample is annotated with a tumor or normal label. For tumor samples, the dataset further annotates the cancer type and ensures that the annotated cancer type is paired with the tissue of cancer origin. At the single-cell resolution, the pan-cancer dataset also annotates each cell with the corrected copy number variation inference results.

#### OOD dataset

We collected a brand-new dataset^49^ as the out-of-distribution (OOD) dataset to evaluate the generalization ability of the model. Although the pan-cancer dataset includes a variety of measurement techniques, in order to further verify the out-of-distribution generalization ability, the validation dataset we need should come from new measurement techniques. The OOD dataset contained 400 thousand cells and 1,500 tissue samples from 37 breast cancer (BRCA) patients. In this dataset, all samples were malignant breast cancer, so all bags were assigned the BRCA label. At the cellular resolution, the dataset has annotated malignant cells and other cell types. For patient samples, the dataset contains information about different diagnostic stages of BRCA.

#### AML dataset

To demonstrate the analysis of new cancer type other than the 30 cancer types in the pan-cancer dataset, we took acute myeloid leukemia (AML) as an example and selected the AML dataset^52^. This dataset contains 35 tumor bags and 6 normal bags, with a total of 30,000 single cells. At the single-cell resolution, the dataset includes annotation information of malignant cells and other cell types, and at the patient sample resolution, it contains the treatment time.

#### Bulk AML dataset

To validate the AML subtype-specific genes obtained from the AML dataset, we used an independent bulk AML RNA-seq dataset^54^ for survival analysis. This dataset measured 671 AML patients and recorded the time when the death events occurred.

### Comparison methods

#### KIDA

KIDA^34^ belongs to the prediction methods based on pseudo-bulk. For a patient sample, the single-cell data is averaged as the bulk expression data. KIDA initializes the parameters based on the pathway annotation, and converts the input bulk gene expression into specific functional patterns. The self-attention mechanism learns the interactions of the functional patterns and outputs the prediction results of the bulk samples.

#### DEGAS

DEGAS^36^ learns the joint latent space of patient samples and single cells simultaneously. It uses the maximum mean discrepancy loss function to reduce the differences between entities at the two resolutions, and then transfers the labels of sample entities to single-cell entities in the joint space. The main computation of DEGAS is concentrated on the integration of patient sample and single-cell data.

#### scIDST

scIDST^37^ belongs to instance-based MIL. All instances in a bag are annotated with the label of the bag as pseudo-labels. scIDST sets up a fully connected network to learn the relationship between instances and pseudo-labels, and introduces probabilistic labels to mitigate the possible noise in the instance pseudo-labels.

#### scVI+MIL

This is an implementation of bag-based MIL. First, scVI^58^ is used for unsupervised learning of instance-level representations of single-cell data. Then, the standard bag-based multi-instance learning process is employed to learn the relationship between instance representations and bag labels. For bag-based multi-instance learning, atention pooling is adopted, and the implementation in MultiMIL^38^ is used instead.

### Application

#### Identification of target cell cluster-specific genes

In the single-cell resolution atlas, for any given instance, the instance encoder of scTransMIL returns the attention of the last transformer layer. We calculate the average for all heads to obtain the gene attention score specific to the current instance. Further, we collect all the instances in the target cell cluster and calculate the average attention of these instances. We rank the genes according to the attention scores and select the top 100 genes as the genes specific to the target cell cluster. In this study, we set two groups of target cell clusters respectively: the tumor cell cluster of the specified cell type and the normal cell cluster of the specified cell type.

#### Differential expression analysis

Differential analysis is a comparative method based on attention inference. In this study, we set up two differential analysis methods. For the first method, we randomly selected a malignant sample and performed differential analysis by comparing the malignant cells and normal cells in this sample. For the second method, we selected all the samples from patients in the pan-cancer dataset and conducted differential analysis by comparing the malignant cells and normal cells in these samples. The differential analysis followed the standard procedure of scanpy^59^.

#### Cancer subtype-specific genes analysis

We used an independent bulk dataset to validate the cancer subtype-specific genes inferred by the attention mechanism of scTransMIL. First, we defined the genes specific to each cancer subtype based on attention results, with 100 specific genes (gene set) for each subtype. In the bulk data, we selected the samples with a relatively high average expression level of a certain gene set as the samples related to the current cancer subtype. Furthermore, we conducted a Kaplan-Meier survival analysis based on the stratification results of the samples in the bulk data and the recorded occurrence time of death events for each sample.

## Data Availability

All raw datasets used in this study were already published and were obtained from public data repositories. Pan-cancer dataset is available at https://cellatlas.kaist.ac.kr/ecosystem/. OOD dataset is available at https://cellxgene.cziscience.com/e/34f5307e-7b4d-4a48-b68f-2ba844c6414b.cxg/. AML dataset is available at GSE116256 [https://www.ncbi.nlm.nih.gov/geo/query/acc.cgi?acc=GSE116256]. Bulk AML dataset for survival analysis validation is available at https://www.ncbi.nlm.nih.gov/projects/gap/cgi-bin/study.cgi?study_id=phs001657.v2.p1.

## Acknowledgements

This work was supported by the National Natural Science Foundation of China (Grant No. 62176272, to C.Y.-C.C.), Research and Development Program of Guangzhou Science and Technology Bureau (No. 2023B01J1016, to C.Y.-C.C.), and Key-Area Research and Development Program of Guangdong Province (No. 2020B1111100001, to C.Y.-C.C.). Fan Yang was supported by the Young Elite Scientists Sponsorship Program by CAST (2023QNRC001).

## Author Contributions

F.Y. and J.Y. conceived and designed the project. Z.T. and F.W. developed the algorithm and conducted experiments under the supervision of J.Y.. Z.T. and F.W. analysed the results. Z.T. and F.W. wrote the manuscript. Z.T. finished the figures under the guidance of F.W. and J.Y.. J.Y., J.S. and C.Y.-C.C. gave suggestions for the manuscript. F.W. and F.Y. gave suggestions for the project. Z.T. and F.W. contributed equally. All authors reviewed and approved the manuscript.

